# Strong Episodic Selection for Natural Competence for Transformation Due to Host-Pathogen Dynamics

**DOI:** 10.1101/275651

**Authors:** Nathan D. Palmer, Reed A. Cartwright

**Affiliations:** Division of Biological Sciences, University of California, San Diego, La Jolla, CA 92093 USA; Barrett, The Honors College, Arizona State University, Tempe, AZ 85287 USA; The Biodesign Institute and School of Life Sciences, Arizona State University, Tempe, AZ 85287 USA

**Keywords:** coevolution, recombination, red queen, phage, bacteria, Lotka-Volterra

## Abstract

Sexual recombination only occurs in eukaryotes; however, many bacteria can actively recombine with environmental DNA. This behavior, referred to as transformation, has been described in many species from diverse taxonomic backgrounds. Transformation is hypothesized to carry some selective advantages similar to those postulated for meiotic sex in eukaryotes. However, the accumulation of loss-of-function alleles at transformation loci and an increased mutational load from recombining with DNA from dead cells create additional costs to transformation. These costs have been shown to outweigh many of the benefits of recombination under a variety of likely parameters. We investigate an additional proposed benefit of sexual recombination, the Red Queen hypothesis, as it relates to bacterial transformation. Here we describe a model showing that host-pathogen coevolution may provide a large selective benefit to transformation and allow transforming cells to invade an environment dominated by otherwise equal non-transformers. Furthermore, we observe that host-pathogen dynamics cause the selection pressure on transformation to vary extensively in time, potentially explaining the tight regulation and wide variety of rates observed in naturally competent bacteria. Host-pathogen dynamics may explain the evolution and maintenance of natural competence despite its associated costs.

## 1 Introduction

Many bacterial species actively import DNA into their cells from the environment and incorporate it in the their genomes. This process, which is roughly similar to sexual recombination, is known as transformation and is foundational to modern molecular biology. However, the evolutionary explanation for this important phenomenon remains unclear despite extensive study [1, 2]. It is well known that transformation plays a key role in the evolution of species which undergo it, distributing allelic variation throughout populations, disrupting linkage disequilibrium, and even introducing novel genes to the species [3, 4, 5, 6]. These consequences of transformation carry a host of potential benefits and risks, the balance of which may change across circumstances. The complexity in the evolutionary effects of transformation has resulted in many unanswered or incompletely answered questions. For example, what conditions may select for or against transformation? To what extent is the selective pressure dependent on the rate of transformation? Which of these conditions are likely to be relevant to the last competent ancestor? Which are relevant to the maintenance of the trait today?

The potential benefits of natural competence are threefold: incoming DNA can be degraded to provide nucleotides for DNA replication [7], serve as a template for repairing a damaged genome [8], or provide new allelic combinations that may increase fitness [1, 9].

The nutritional benefit of taking up and degrading DNA for its nucleotide monomers is clearly apparent, and some have suggested that transformation may be a side effect of this behavior, even suggesting that the nutritional effect alone could be sufficient to maintain competence [7]. This idea is supported by the fact that starvation activates competence in *B. subtilis* [10], and the Sxy transcription factor, a master regulator of competence in several gram negative species, is regulated by intracellular purine levels [11]. However, the nutrition hypothesis falls short of explaining several mechanistic components of the transformation system, including the conserved proteins DprA, SsbB, and DpnA, which act to protect DNA imported through ComEC from degradation by intracellular nucleases [2]. It also fails to explain why bacteria in DNA-rich environments, such as the respiratory pathogen *S. pneumoniae*, regulate both competence expression and DNA secretion via complex mechanisms such as quorum sensing [12]. Additionally, other species, such as *V. cholerae*, have been shown to turn off genes involved in DNA catabolism, such as extracellular nucleases, upon competence induction [13].

Another idea is that bacteria may take up foreign DNA to serve as a template for homologous repair of double strand DNA breaks. The initial lines of evidence for this hypothesis were based on increased survival of transforming types upon experimental challenge with DNA damaging agents [8] and the observation of regulatory schemes consistent with the repair hypothesis in some species of competent bacteria [14]. More recent research has led to the suggestion that competence may serve as a sort of replacement for the SOS response system in *P. pneumoniae* and *L. pneumophila* [15]. Nevertheless, this explanation makes little sense when applied to other species such as *B. subtilis*, which contains an intact SOS system, and *S. thermophilus*, in which competence is regulated in opposition to the SOS system [2].

Much of the evidence used to argue for the nutrition and repair hypotheses relies on analysis of the regulatory schemes by which competence is induced. However, research over the last 20 years has demonstrated that the complex regulatory mechanisms governing the expression of competence genes, involving alternative σ-factors, quorum sensing, and multiple levels of specific and global transcription factors are inconsistent with explanation by either of these hypotheses alone. Furthermore, both the biochemical means of regulation and the environmental cues that induce transformation vary widely and appear to have evolved independently among competent species [2, 13]. Given these findings, the environmental cues and regulatory signals on a trait in a few species is not sufficient to broadly infer the evolutionary history of that trait.

The controversy over the evolutionary story behind natural transformation in bacteria has continued for over two decades now and seems little closer to being resolved. The large amount of new and more diverse data has clarified the situation some, but much uncertainty remains. For a more in-depth discussion of the evolution of natural competence, see [1], [2], and [13].

Transformation by homologous gene recombination in bacteria is thought to provide some of the same benefits as meiotic sex in eukaryotes, such as slowing the rate of accumulation of deleterious mutations and combining beneficial alleles from different lineages [16, 17, 18]. However, transformation has additional costs compared to meiotic sex. Cells asymmetrically lose competence by transforming in loss-of-function alleles at loci necessary for competence in mixed populations [19]. Furthermore, although there is evidence that some populations engage in autolysis and/or DNA secretion for purposes of transformation [20], the pool of homologous free DNA is likely to be largely composed of DNA from dead cells for most competent species, and therefore loaded with more deleterious mutations than the living population (bad genes effect) [19, 21]. Despite these costs, laboratory evolution experiments have suggested that transformation may speed up population-level adaptation rates and provide a selective advantage due mostly to population genetic consequences rather than nutrition or repair [22].

Models of the evolution of recombination under mutation/selection dynamics are highly dependent on opaque parameters, especially the distribution of fitness effects of new mutations [23, 24]; however, computational models have shown that asymmetrical loss and the bad genes effect outweigh the single population benefits of transformation (Hill-Robertson Effect and breaking Muller’s ratchet) under a wide variety of likely conditions [19, 21].

Another proposed benefit of meiotic sex is increased adaptability in arms races between hosts and pathogens or predators and prey (Red Queen hypothesis) [25]. It is well known that host-pathogen coevolution is a major factor in the evolution of some bacteria-phage systems [26], and it has been speculated to exert some selective pressures on the homologous recombination rate [1], but to our knowledge this idea has not been rigorously investigated in either computational models or experimental work. The large variations in selective pressures created by host-pathogen coevolution also fit previous models of transformation evolution and maintenance based on short, periodic bursts of strong selection for transformers [27]. Given the widespread presence of homologous recombination by transformation in bacteria, combined with the difficulty in describing a convincing advantage to the phenotype based on traditional mutation and selection evolution, we have hypothesized that host-pathogen dynamics contribute to the evolution and maintenance of natural competence for transformation in bacteria.

To investigate this hypothesis, we have developed a series of computer models with which we can analyze the consequences of transformation in a coevolving population across different environmental and genetic conditions. Using these models, we demonstrate that large cyclic variations in population size due to phage predation select for transformation, allowing a very small subset of transformers to invade a large, established population of non-transformers. We also find evidence for negative frequencydependent selection on transformation, which may provide insight into competence-controlling mechanisms such as stochastic switching in *B. subtilis* [28] and coordinated competence in *S. pneumoniae* [12].

## 2 Methods

### 2.1 Deterministic Model

We have modeled bacterial growth in the presence of phages with a modified form of Lotka-Volterra differential equations [29]. Our model incorporates one marker locus and one transformation locus, each with two alleles. The marker locus determines the susceptibility of the bacteria to phage predation, DNA uptake and homologous recombination, and a cell may be either competent ‘*t*’ or non-competent e.g. a cell-surface protein that the phage recognizes to bind and infect the bacterium. There are two alleles at this locus, which we call ‘1’ and ‘2’. The transformation locus represents the genes needed for ‘*n*’ at this locus. There are two phage types, each corresponding to one of the marker alleles in the bacterial population. When a bacterium is lysed by a phage, it releases its alleles into the environment individually. These free alleles can be taken up and incorporated, replacing their current allele, by living bacteria if they are competent at the transformation locus.

We keep track of 10 populations in the model: 4 bacterial types, *B*_1*t*_, *B*_2*t*_ (transforming), *B*_1*n*_, *B*_2*n*_ (non-transforming); 2 phage types, *P*_1_, *P*_2_; and 4 DNA types, *D*_1_, *D*_2_, *D*_*t*_, *D*_*n*_. These populations change with time according to Equations 1, where *α* is the rate of cell division, *β* is the rate of phage infection, *γ* is the rate of phage decay, *δ* is the phage fertility rate, *ρ* is the transformation rate, *ζ* is the rate of free DNA decay, and *κ* is the bacterial carrying capacity of the environment. *N* is the sum bacterial population size. Parameters in this model were chosen in accordance with experimental work analyzing bacteriophage coevolution [30].

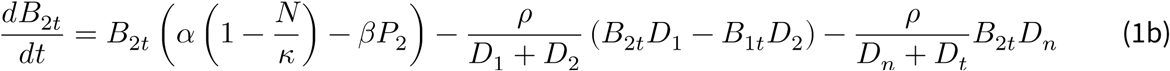

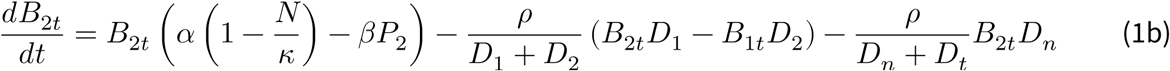

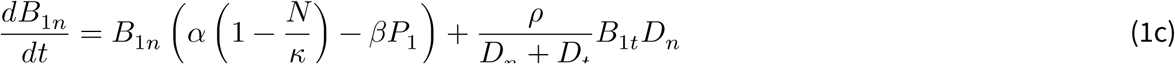

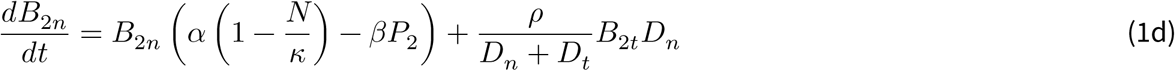

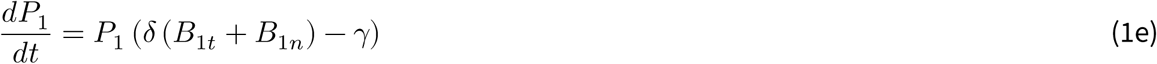

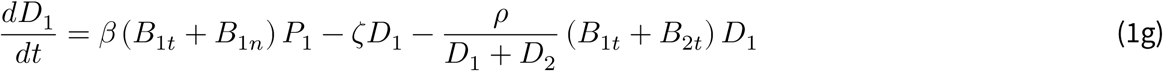

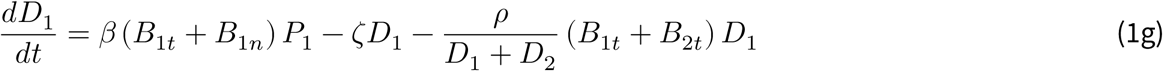

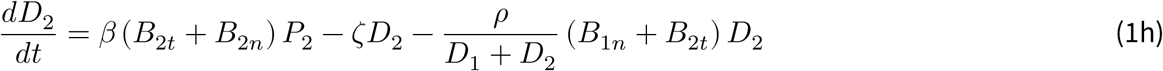

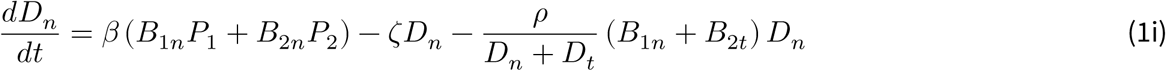

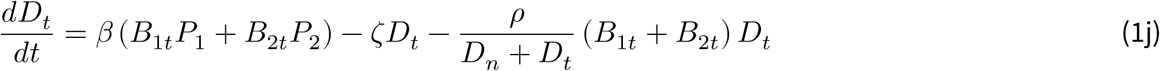

Boom-bust cycles in Lotka-Volterra dynamics without transformation (*ρ* = 0) can be described by enclosed curves in phase space by solving the system of equations for 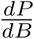 for all values of *t*. Integrating the log of this expression for each allele (allele 1 shown here, 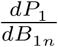), results in a conserved, constant quantity, *V*.

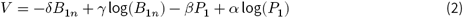

*V* can be thought of as a measure of displacement of the population from equilibrium. It is maximized at equilibrium, and as *V* decreases, the magnitude of the fluctuations in population size increases.Note that without transformation, *V* remains constant over time. However, because transformers can switch types and escape predation, when the transformation rate *ρ* > 0, *V* becomes a function of time.

We used *V* to investigate what conditions make a population susceptible to transformer invasion. We initialized simulations of displacement from equilibrium by manipulating the initial *V*, then setting *B*_1*n*_ and *B*_2*n*_ at the minimum and maximum, respectively, on the curves described by each *V*. We implemented this model in R [31] using the gsl [32] and deSolve [33] packages.

### 2.2 Stochastic Model

In the deterministic model, a population at equilibrium will never leave it. However, in reality, stochasticity would cause fluctuations in the population sizes, naturally creating boom and bust cycles. We used the Gillespie algorithm to implement the model in fully-stochastic time to determine if stochasticity can induce the cyclic conditions that allow transformers to invade [34].

The stochastic model is derived from the differential equation model, but presented in an event-based paradigm.

| Event | Rate |
| --- | --- |
| Cell division | $\alpha B_x$ |
| Lysis and phage replication | $\beta B_x P_x$ |
| Transformation with new allele | $\frac{\rho}{\sum_i D_i} B_x D_x$ |
| Phage decay | $\gamma P_x$ |
| Free DNA decay | $\zeta D_x$ |
| Immigration | $\lambda$ |
| Phage mutation | $\epsilon$ |

Unlike the deterministic model, populations here are susceptible to extiction due to discrete population sizes. To protect against extinction, we incorporated slow mutation from one phage type to the other and steady migration of all bacterial types into the model.

At a given point in the simulation, all rates are calculated based on the current state of the population, and the time until the next event is drawn from an exponential distribution with rate equal to the sum of the rates of all events. At this time, the event that occurs is drawn based on its relative probability and executed. This is repeated until termination criteria are fulfilled. Parameters in the stochastic model were scaled down from the more biologically realistic parameters in the deterministic model to make the model computationally tractable. We implemented this model in R [31] using the GillespieSSA [35] package.

## 3 Results

### 3.1 Deterministic Model

We have modeled a coevolving population of bacteria and phages to examine the selective pressures on homologous recombination by transformation. Transforming bacteria are initially very rare, but invade and become dominant over time (Figures 1, 2). Invasion proceeds when the cycles between the two infection alleles are opposed. When levels of *B*_1*t*_ and *B*_1*n*_ are high and about to crash, many of the *B*_1*t*_ transform to *B*_2*t*_,which are at low numbers but about to grow as phage *P*_2_ decay outpaces reproduction due to low numbers of hosts. This allows the proportional growth rate of *B*_2*t*_, (*αB*_2*t*_ + *ρB*_1*t*_*D*_*2*_), to exceed that of *B*_2*n*_, (*αB*_2*n*_), during the growth phase.

**Figure 1:**
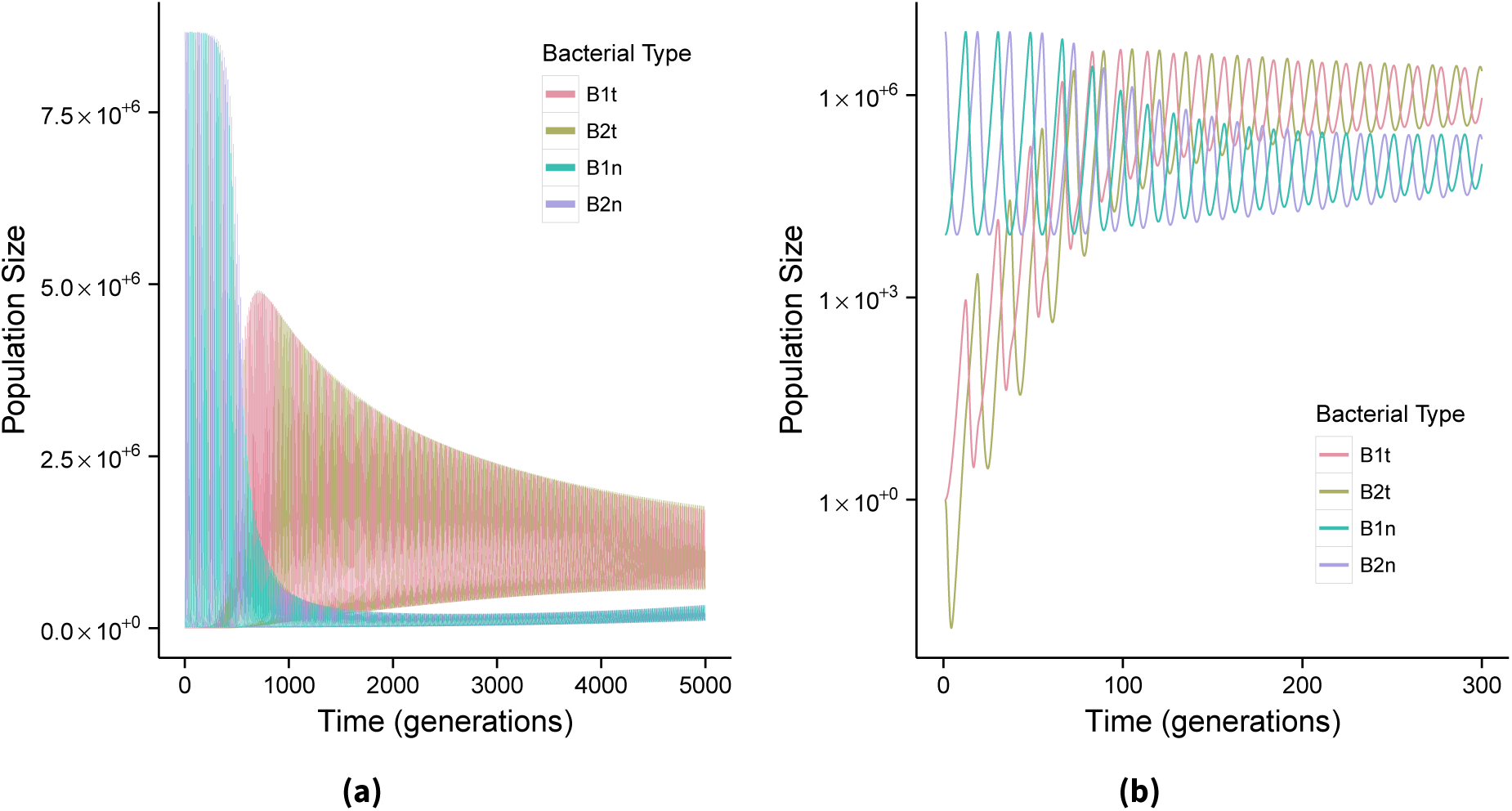
Simulation dynamics in the deterministic model shown at normal **(a)**, and log **(b)** scale.Initial transformer frequency = 10^−6^, *α* = 1, *β* = 1×10^−7^, *γ* = 0.25, *δ* = 2 × 10^−7^, *ζ* = 0.1, *κ* = 10^13^, **(a)**: *ρ* = 5 × 10^−4^, **(b)**: *ρ* = 5 × 10^−3^.

**Figure 2:**
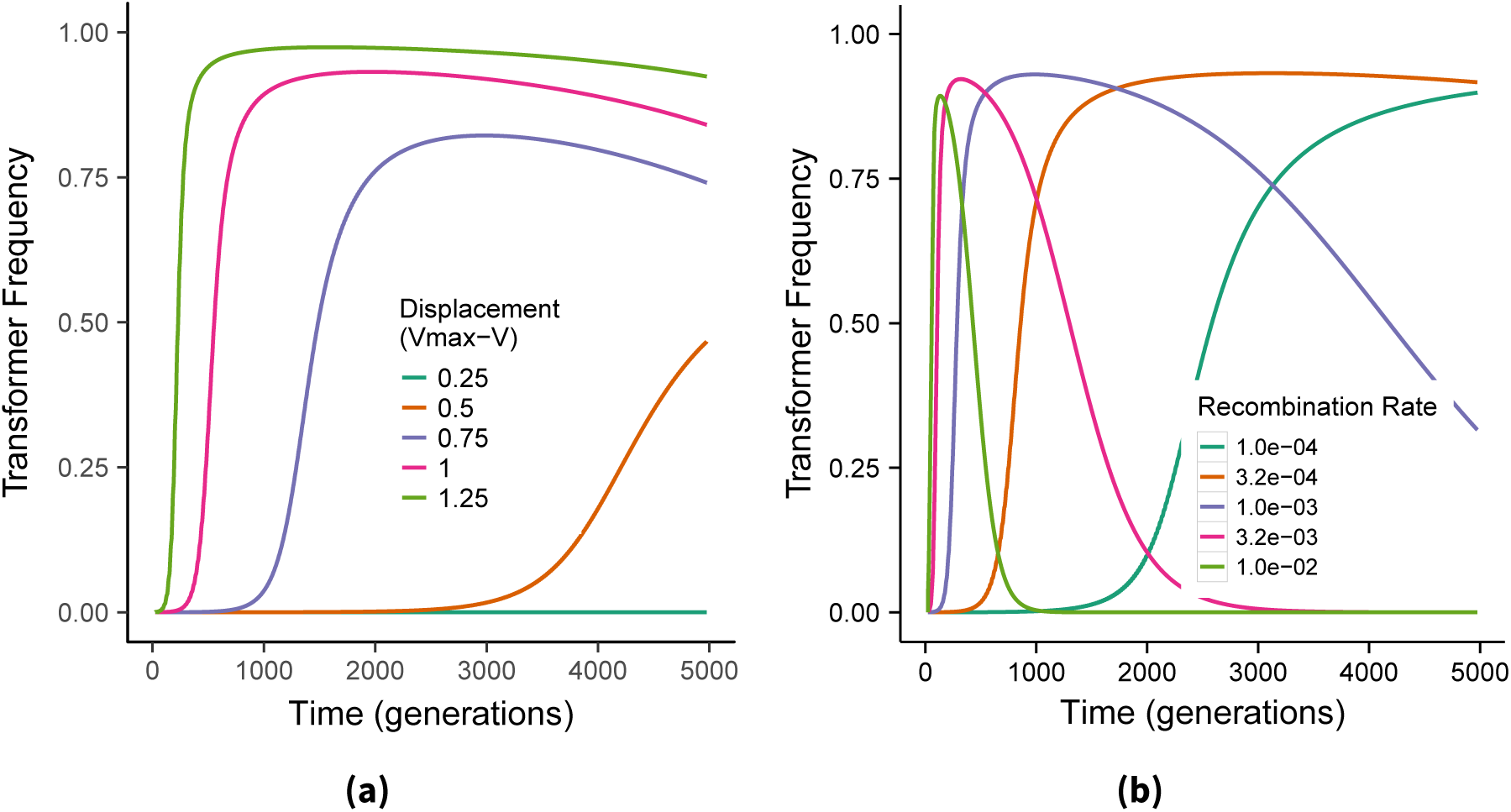
Transformers invade a population of non-transformers under Lotka-Volterra dynamics dependent on recombination rate and amplitude of predation cycles. **(a)**: Transformers invade faster, to a higher maximum frequency, and persist longer when predation cycles are large. Invasion is abolished when cycles are sufficiently small. Initial transformer frequency =10^−6^, *α* = 1, *β* = 1 × 10^−7^, *γ* = 0.25, *δ* = 2 × 10^−7^, *ρ* = 5 × 10^−4^, *ζ* = 0.1, *κ* = 10^13^. **(b)**: Transformer invasion occurs faster but is less stable with increasing recombination rate. Same simulation parameters as **(a)**, Δ*V* = 1.0

To better visualize this difference, we have unpacked the components of the growth rate of *B*_1*n*_ during invasion in Figure 3. Because of new transformants coming from type *B*_2*t*_, the *B*_1*t*_ type stops decreasing and begins growing earlier in the phage cycle and grows faster during that period of early growth.This continues until type *B*_2*t*_ is no longer abundant, resulting in a decrease in transformation and a reversion of the *B*_1*t*_ growth rate to approximating the base rate not including transformation. This phenomenon repeats each cycle, with magnitude growing logarithmically each time, resulting in the invasion pattern seen in Figure 1b.

**Figure 3:**
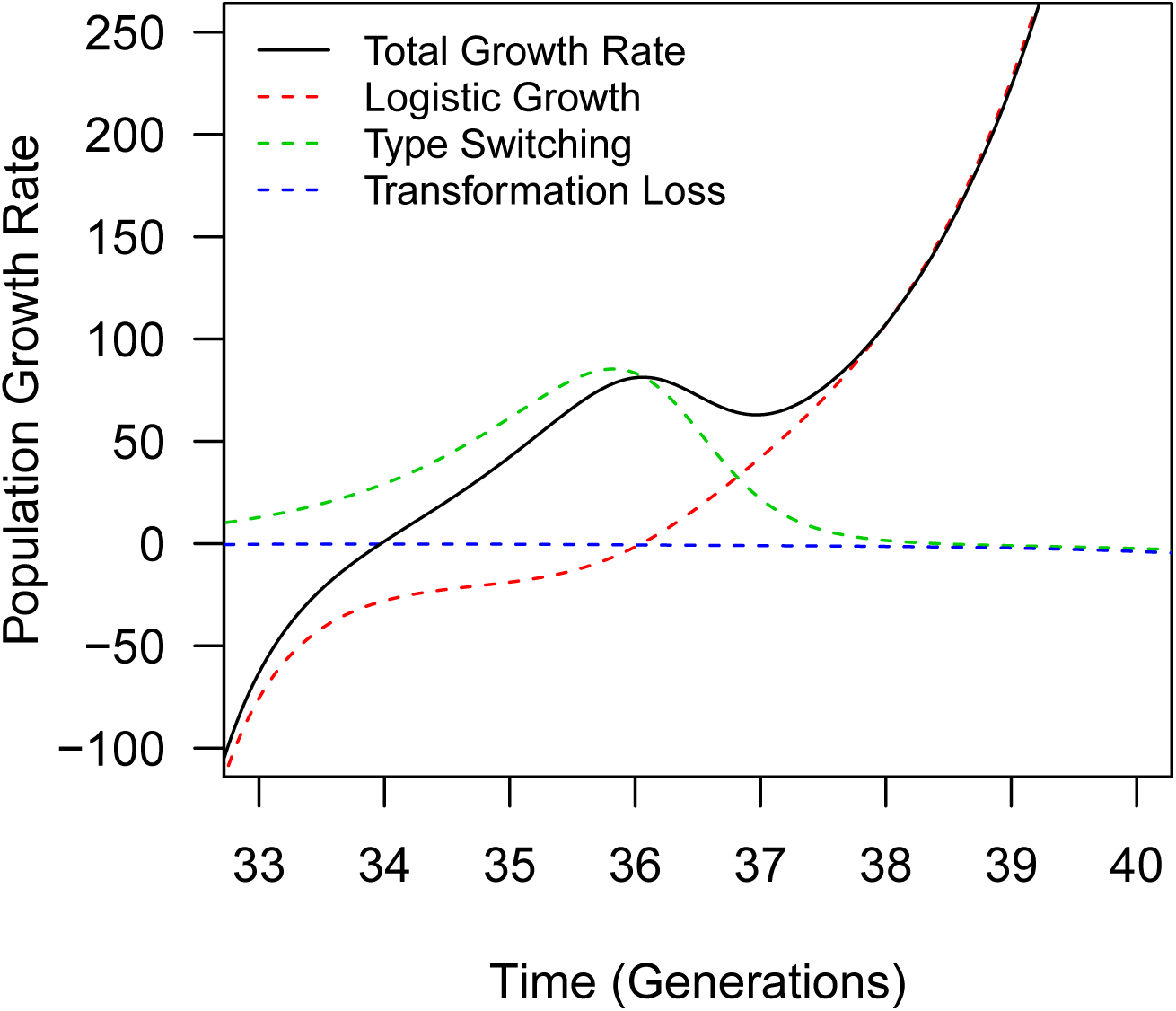
Growth rate components of transforming bacterial type *B*_1*t*_ during invasion. The solid black line represents the total growth rate; dashed lines are the three main components of equation 1a. The red dashed line represents logistic growth and phage predation, the green dashed line is type switching between types *B*_1*t*_ and *B*_2*t*_ by transformation, and the blue dashed line is type switching from *B*_1*t*_ to *B*_1*n*_ by transformation.

This effect, compounded over several cycles, allows the transformers to invade. However, when the transformers are the dominant type in the population, this phage escape feeds back into the phage population growth and death rate, smoothing the boom/bust cycles. When these cycles become small, the transformers are no longer able to overcome the asymmetrical loss of transformation alleles, and non-transformers gradually re-invade (Figure 1).

The term which represents the difference in growth rate between a transformer, say *B*_2*t*_, and the corresponding non-transformer, *B*_2*n*_, is *ρB*_1*t*_*D*_2_. This term, and therefore transformer invasion depends on (1) the transformation rate, *ρ*, and (2) the population sizes of transformers of type 1 and free DNA of type 2. Both of these last terms are directly affected by the magnitude of boom and bust cycles.

Transformers are only able to invade when population fluctuations due to host-pathogen dynamics are sufficiently large. The maximum transformer frequency and length of invasion also depend on this factor (Figure 2a). Since transformers invade by taking advantage of alternating selective pressures due to phage boom and bust cycles, it is expected that the magnitude of these cycles would determine the magnitude of transformer advantage. The recombination rate also affects the speed of transformer invasion (Figure 2b). Rapid recombination leads to quick invasion, but also makes the population more susceptible to reinvasion due to faster loss of transformation alleles, while lower recombination rates slow both invasion and reinvasion.

Under our model, the rate at which bacteria carrying the transformation allele invade is inversely proportional to the magnitude of population fluctuations. The speed of invasion asymptotically slows approaching a threshold value defining how far a non-recombining population must venture from equilibrium to be susceptible to transformer invasion (Figure 4a). The length of invasion shows a similar pattern of dependence on the recombination rate (Figure 4b). This equilibrium value depends on the recombination rate of the transformers: increasing recombination by transformation pushes the threshold closer to equilibrium.

**Figure 4:**
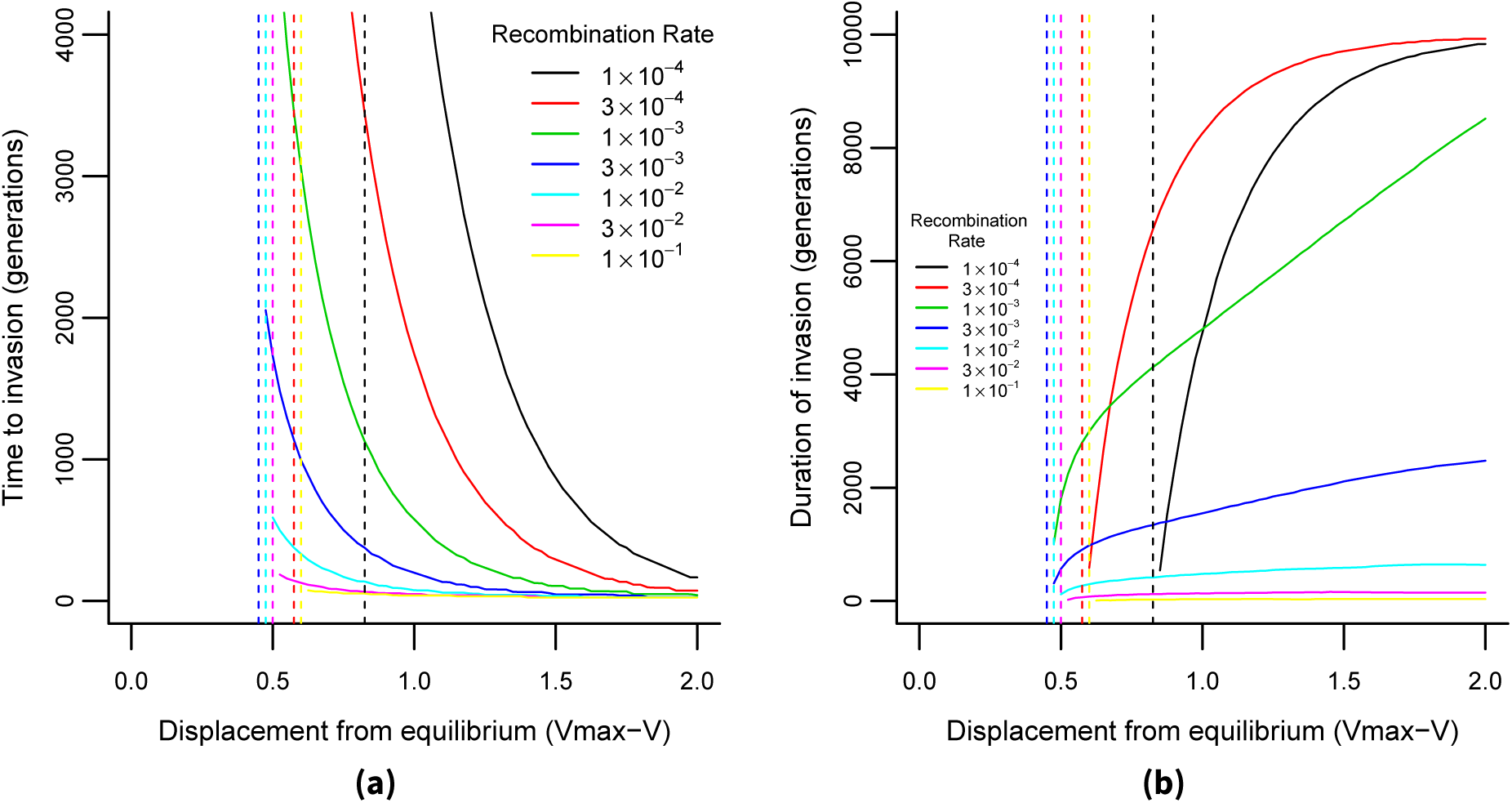
**(a)**: Invasion speed decreases with predation cycle size, asymptotically approaching a threshold size, below which transformers cannot invade. The threshold size depends on the recombination rate. The invasion threshold is minimized at recombination rate ≈ 3 × 10^−3^. **(b)**: After invasion, transformers decline due to asymmetrical loss of transformation alleles with a rate proportional to recombination. Parameters are the same as in Figure 2.

### 3.2 Stochastic Model

We initialized simulations containing only non-transformers at equilibrium and let them burn-in for 100 generations. Two transformers of each type (*B*_1*t*_ and *B*_2*t*_) were added to the simulation. The transformers became the dominant group in the population, defined as *B*_1*t*_ > *B*_1*n*_ and *B*_2*t*_ > *B*_2*n*_, with a frequency dependent on the recombination rate. Probability of invasion increases with increasing recombination rate until approximately 3 × 10^−2^, where probability of invasion is maximized (≈ 4 –fold greater than non-transformers). At rates higher than this, the probability of invasion sharply decreases. It is important to note the magnitude of the optimal transformation rate in this model may not correspond to that in nature due to the parameters used; however, the pattern of the distribution should hold. The transformation rate estimates from the deterministic model where experimentally validated parameters were used are more likely to be close to reality.

Unlike the deterministic model, the length of invasion is maximized at the same recombination rate as the frequency or time to invasion. This may be explained by the fact that stochasticity prevents the transformers from reducing the boom/bust cycles through escape to the same extent as in the deterministic model.

This model demonstrates that stochastic effects can naturally create conditions that allow transformers to invade, even when the simulation is initialized at a non-transformer dominant equilibrium (Figure 5). In fact, the stochastic simulations cycle through the dynamics observed in the deterministic model: the dominant type in the population alternates between transformers and non-transformers.

**Figure 5:**
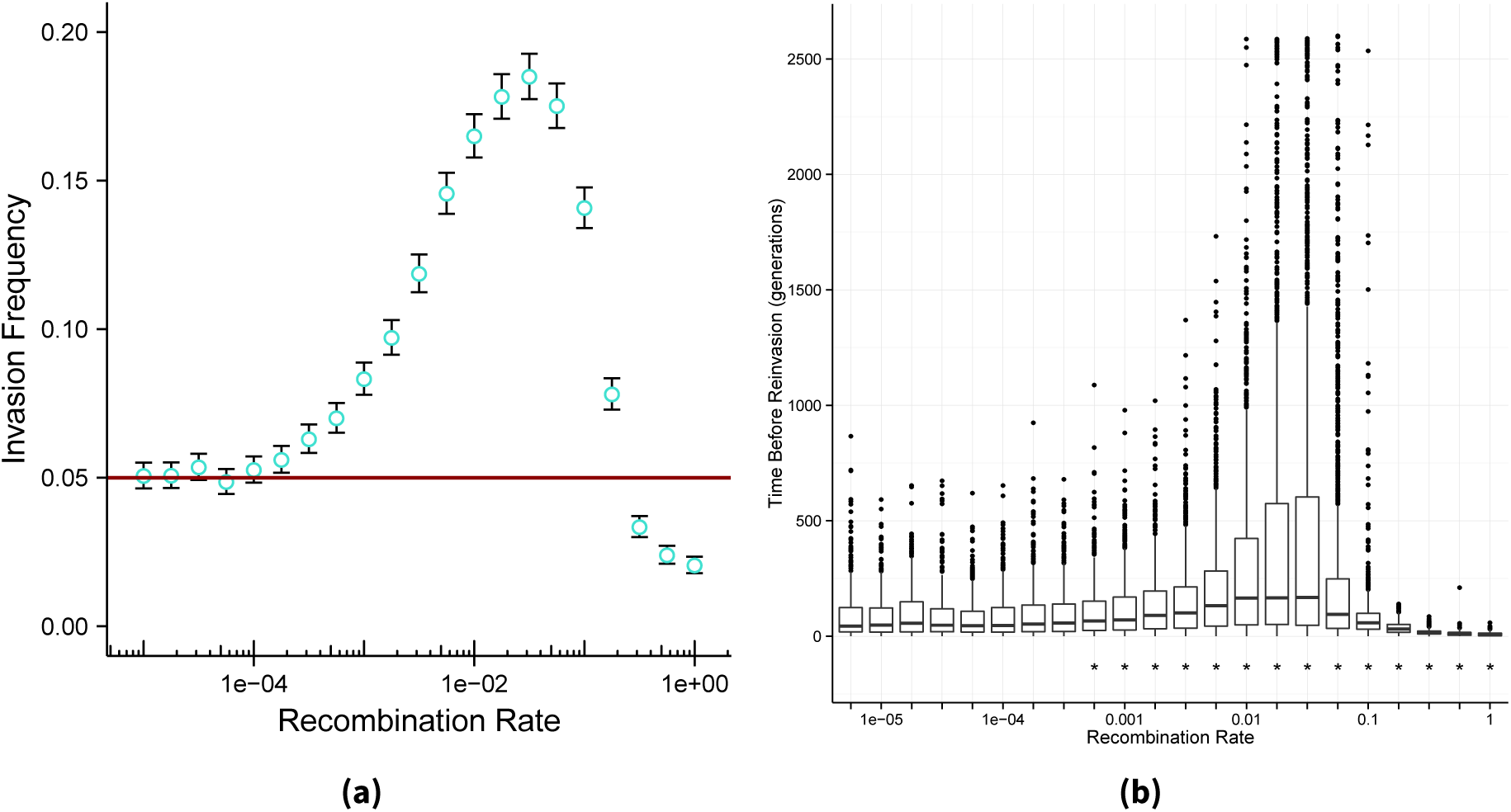
Transformers invade populations starting at equilibrium due to stochastic effects. **(a)**: Frequency of transformer invasion after burn-in. The red line represents invaders with no recombination. Each data point represents 10^5^ independent simulations. **(b)**: Duration of invasions in the simulations in **(a)**. *α* = 0.5, *β* =0.005, *γ* = 0.5, *δ* = 0.01, *ρ* = 0, *ζ* = 0.3, *k* = 10^4^, *λ* = 0.05, *ε* = 0.02)

## 4 Discussion

Our results indicate that host-pathogen dynamics provide a selective advantage to transforming bacteria which largely depends on two factors: the rate of transformation, and the size of population fluctuations. The population fluctuations are determined by the growth rate of the bacteria, the infection rate, the burst size, and the phage decay rate. We estimate that under biologically likely parameters, invasion occurs most frequently at a recombination rate on the order of 3 × 10^−3^. To explain these results, we conceptualize the following model.

In large boom-bust cycles, the crash of one population due to predation opens up a new niche with plentiful resources for a population immune to the previous phage infection. Transformers maintain extra genetic variation in the form of the free DNA pool, which is not subject to selection due to phage predation. These alleles can be reintroduced into the population quickly, allowing the transforming types to have a significant advantage in colonizing the new space. Due to transformation from the type under phage attack, transformers are able to both begin colonizing the new space earlier, and grow faster during the early colonization phase. In the short term, this advantage is paramount, allowing transformers to invade and dominate. In the long run, however, as smaller cyclic variation reduces the transforming advantage, non-transformers re-invade. This is mediated by transformers incorporating free DNA disrupting or eliminating its ability to transform by homologous recombination, while a non-transformer never gains the genes necessary for homologous recombination by transformation [19]. As a result, the allele frequency of functional transformation alleles will gradually decrease in the absence of compensatory selective pressures. For a transformation phenotype to persist, those selective pressures must be sufficiently strong to overcome this effect. Alternatively, since bacteria are under attack by phages, transduction may also play a role in preserving transformation alleles. This can be explored in future work.

Under stochastic variation, cycles will be randomly aligned and misaligned in time. Therefore the selection coefficient of the transformation allele will switch signs frequently and be sharply dependent on the rate of recombination. Theoretical work has predicted that episodic selective sweeps have led to the evolution of competence for transformation [27], and we hypothesize that host-pathogen coevolution provides these sweeps. Indeed, experimental populations of bacteria coevolving with phages have shown these kinds of fluctuating conditions [26, 36].

It is important to note that this model is valid not only for phage predation, but also for predation by larger organisms such as eukaryotic protists or immune cells. Here, we have chosen to base our model on phages, as they are a relatively universal selective problem for bacteria. However, transformation is somewhat common among pathogenic bacteria, e.g. *S. pneumonia, N. gonorrhoeae, H. pylori*, etc. For these species, immune escape may be a more critical selective force. Our model is applicable to those situations as well given the partial symmetry in the Lotka-Volterra model [29, 37]. It would be a judicious test of our model to experimentally set parameters to the environment of a variety of species, and determine if the model accurately predicts their different recombination rates.

We have shown that host-pathogen dynamics can provide a mechanism for transformation alleles to become prevalent in a population. However, the strong episodes of selection which underlie that mechanism are interspersed between relatively longer periods in which transformation is maladaptive. The persistence of the trait may be reconciled with this selective regime by considering a meta-population, spatially explicit model of evolution.

New research is suggesting that the evolution of many bacterial traits, including natural competence for transformation is best thought of in spatially explicit terms [38]. A complex spatial structure may result in a situation in which any given population in a given space has only infrequent conditions selecting for transformation, such as those encountered in colonization of new territory after localized extinction or near-extinction, but transformers can always find their niche somewhere, even if it is frequently in motion. Complex regulatory schemes like those observed in nearly all competent species [2], may further help transforming bacteria navigate this evolutionary landscape. New experimental approaches using spatially controlled bacterial populations may further validate this exciting idea.

## 6 Acknowledgments

Christian Sievert and Xuan Wang provided helpful feedback on this manuscript.

